# Towards Wearable Electromyography for Personalized Musculoskeletal Trunk Models using an Inverse Synergy-based Approach

**DOI:** 10.1101/2024.07.23.603973

**Authors:** Jan Willem A. Rook, Massimo Sartori, Mohamed Irfan Refai

## Abstract

Electromyography (EMG)-driven musculoskeletal models (EMS) of the trunk are used for estimating lumbosacral joint moments and compressive loads during lifting tasks. These models provide personalized estimates of the parameters using information from many sensors. However, to advance technology from labs to workplaces, there is a need for sensor reduction to improve wearability and applicability. Therefore we introduce an EMG sensor reduction approach based on inverse synergy extrapolation, to reconstruct unmeasured EMG signals for different box-lifting techniques. 12 participants performed an array of tasks (squat, stoop, unilateral twist and bilateral twist) with different weights (0 kg, 7.5 kg and 15 kg). We found that two synergies were sufficient to explain the different lifting tasks (median variance accounted for of 0.91). Building upon this, we used two sensors at optimal subject-specific muscle locations to reconstruct the EMG of four unmeasured channels. Evaluation of the reconstructed and reference EMG showed median coefficients of determination (*R*^2^) between 0.70 and 0.86, with median root mean squared errors (RMSE) ranging from 0.02 to 0.04 relative to maximal voluntary contraction. This indicates that our proposed method shows promise for sensor reduction for driving a trunk EMS for ambulatory biomechanical risk assessment in occupational settings and exoskeleton control.

## I. INTRODUCTION

In 2020, the global prevalence of low back pain exceeded half a billion cases, with 22% of years lived with disability attributing to occupational factors [1]. Workers who perform occupational tasks such as trunk flexion, rotation, and lifting have an increased risk of injuries [2]–[4]. These injuries result from increased mechanical stress on structures such as intervertebral disks, facet joints, and surrounding tissues, leading to low back pain (LBP) [5]. Compressive spine loads are therefore a predictor for LBP [6]. These loads are generally highest in the caudal end of the lumbar spine [7]. Measuring the lumbosacral joint compressive loads in situ is invasive. There are several modeling approaches to estimate these parameters [8], [9]. However, these approaches do not include measured muscle activations during the lifting tasks, and rather estimate them from the kinematic and kinetic data. In our previous work, we developed an electromyography (EMG)-driven musculoskeletal model (EMS) of the trunk to estimate lumbosacral flexion and extension moments [10]. The approach derives muscle activations from measured EMG and transforms it non-linearly to muscle force along with information on joint position [11]. Thus, the EMS transforms activation signals and joint angles via a muscle contraction dynamics model into muscle forces and joint moments [12]. This model showed a high correlation compared to reference moments estimated using inverse dynamics (ID) (*R*^2^ mean range: 0.88 - 0.94, RMSE: 0.21 – 0.38 Nm/kg). The model was further validated for estimations of compressive loads [13].

The EMS has advantages over traditional ID methods. The ID approach calculates joint moments from body positions and external forces, which requires motion-capture cameras and force plates that are typically available in laboratory settings. Therefore, ID is not suitable for ambulatory assessment of lumbosacral moments [14]. However, an EMS can be driven using information from wearable sensors [13].

The EMS still requires the application of several wearable sensors on the users during the dynamic tasks. This translates to a need for expertise, extended preparation time, and user discomfort [15]. In our previous work, we used 12 EMG sensors and eight inertial measurement unit (IMU) sensors for the EMS [10], to estimate the muscle activity and joint angles respectively. Thus, to improve the wearability of EMG-driven musculoskeletal models, there is a need for sensor reduction for minimal and robust sensor setup with easy application and calibration [16].

Utilizing muscle synergies during dynamic tasks can help reduce the dimensionality of the EMG sensors being used. Synergies describe how the human central nervous system (CNS) coordinates the activation of muscles during specific dynamic tasks [17]. Synergy analysis involves decomposing EMG signals from a group of collaborating muscles into a low-dimensional set of time-independent synergy vectors and time-dependent synergy excitations. Posteriori, matrix decomposition methods, such as non-negative matrix factorization (NMF) can extract these synergies [18]. Muscle activity during movement of the trunk is highly redundant, with the number of synergies usually fewer than the total muscles involved [19]–[21]. For example, Tan et al. [19] showed that three synergies were sufficient to reconstruct EMG signals from eight muscles during stoop lifting. These synergies were time-invariant, which was also found by Sedaghat-Nejad et al. [20] for isometric trunk exertions in 12 directions with equal 30-degree angular intervals in the transverse plane. Wang et al. [21] used high-density EMG with 120 channels and found that only two muscle synergies could explain over 90% of recorded muscle activities in five spinal motions (flexion/extension, lateral bending, axial rotation, sitting, and standing) for healthy subjects.

Reducing dimensionality through synergies offers the potential to minimize the number of sensors required for driving the EMS. One such technique is Synergy Extrapolation (SynX) which facilitates finding synergies based on a multi-objective optimization [22]–[24]. Here, the cost function aims to minimize joint moment tracking errors from ID and unmeasured and residual activation magnitudes [22], [23]. Simultaneously, an EMS can be calibrated to model biomechanical muscle behavior. SynX showed reliable and efficient estimates of unmeasured muscle activation during treadmill walking. Thamid et al. [24] also used the SynX framework to estimate single unmeasured muscle activation for upper extremity tasks. Incorporating the biomechanical behavior of muscles through an EMG-driven musculoskeletal model offers advantages. For example, it enables estimating EMG channels for deep muscles that are difficult to measure. So far, the SynX approach has been used to estimate one muscle [22], [24] or two muscles [23] that were left unmeasured. Bianco et al. [25] estimated 16 EMG signals from eight measured channels during treadmill walking. They initially applied NMF to extract synergy excitations from these eight measured muscles. Subsequently, least squares regression was used to determine the synergy vectors of the 16 unmeasured muscles. The multiplication of these synergy vectors with the measured synergy excitations allowed for the reconstruction of the 16 unmeasured EMG channels. This approach was tested across all possible combinations of eight-muscle subsets, with some subsets achieving over 0.90 variance accounted for (VAF). These approaches highlight the potential of synergy analysis for estimating unmeasured muscles. As our trunk-EMS approach estimates the lumbosacral joint moments and compressive loads with surface EMG robustly [13], here, we explored synergies to improve the wearability of the trunk-EMS [13].

The goal of this study is to assess the feasibility of using synergy extrapolation for estimating the normalized EMG linear envelopes of unmeasured muscles from a reduced sensor set during box lifting. We develop a method for the estimation of synergy excitations during different box-lifting tasks considering several weights. Here, we assume that synergy vectors are subject-specific, and are constant, whereas the synergy excitations can be scaled for different movements and weights. By employing synergy vectors from a prior calibration phase, we aim to reconstruct the missing EMG signals and evaluate their reconstruction accuracy by comparing them to the reference EMG signals.

## II. METHODS

### A. Participants

12 healthy (7/5 M/F; 29.7±12.5 years old, 71.7±9.6 kg and 177.9±8.3 cm tall) participants were included and provided their written consent. None suffered from back injuries. Ethical approval was obtained from the University’s Natural Sciences and Engineering Sciences Ethics Committee (230447).

### B. Experimental Protocol

All participants lifted a box (*w*×*d*×*h* = 40×30×22 cm, empty weight = 1.6 kg) using two symmetric (Stooping (ST) and Squatting (SQ)), and two asymmetric techniques: Unilateral Twisting (UT) and Bilateral Twisting (BT) as shown in fig. 1.

**Fig. 1.**
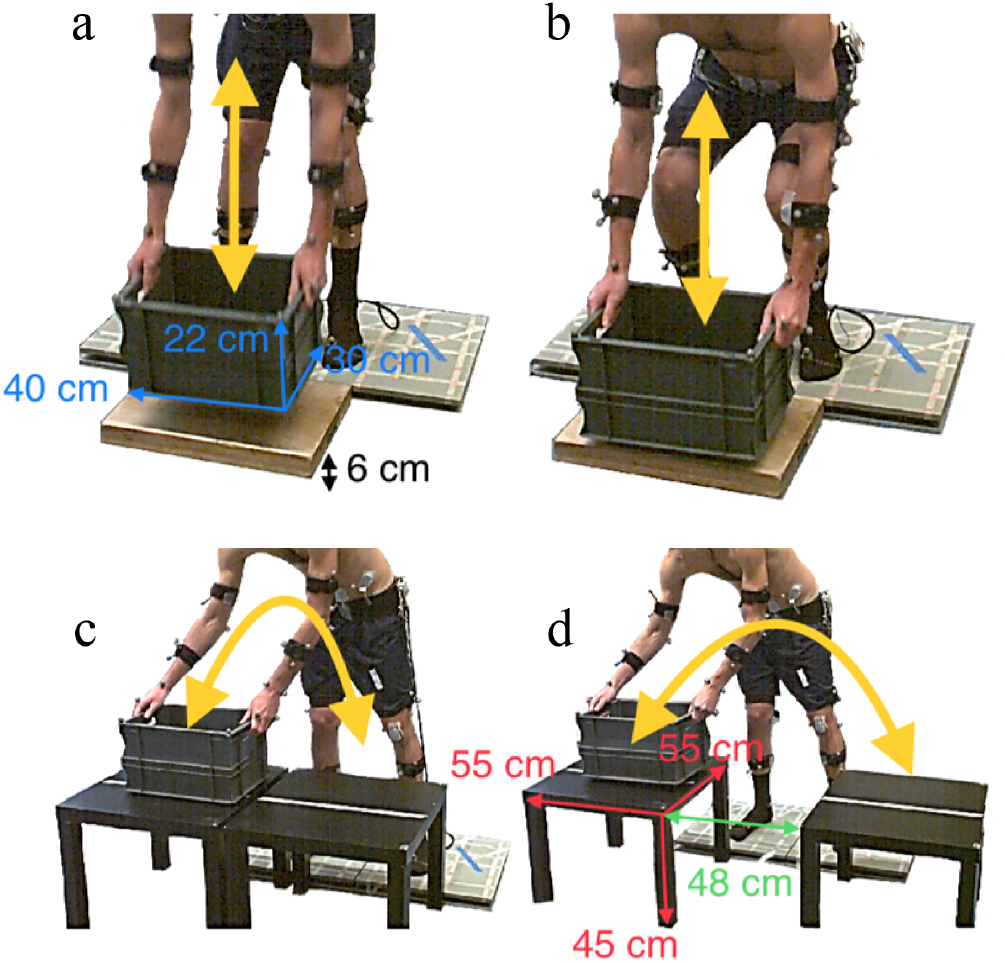
Overview of the four different box lifting techniques: Stoop (a), Squat (b), Unilateral Twist (c), Bilateral Twist (d). Yellow arrows indicate box direction during lifting, blue arrows indicate box dimensions, red arrows indicate table dimensions and the green arrow indicates the distance between the two tables during a bilateral twist.

The ST and SQ techniques involved lifting the box in front of the participant from a wooden platform (*h* = 6 cm). Once the participant was standing upright, the box was placed back in its original position using the same lifting technique in reverse order. Unlike the ST and SQ, the UT and BT techniques involved moving the box from a table (*w*×*d*×*h* = 55×55×45 cm). For UT (fig. 1c), the table was positioned in front of the participant, while for BT (fig. 1d), it was diagonally to the left. The box was then transferred to another identical table diagonally positioned to the right in front of the participant. After standing straight, the participant returned the box to the starting table. These actions include one UT or BT movement. Participants maintained a steady pace by performing all liftings in sync with a 30 beats per minute (bpm) metronome. For each lifting technique, three different weight conditions were considered. The box contained either no additional weight (0 kg), 7.5 or 15 kg^3^. One trial consisted of either six (SQ and ST) or three (UT and BT) repetitions. So the total number of liftings per participant was (6 + 3)(2×3) = 54. Between trials, there was a brief recovery period of approximately 20 seconds for saving data and preparing for the subsequent trial. The trials were grouped for symmetric and asymmetric techniques. The order of symmetric or asymmetric trials, as well as the sequence within these trials, were randomized.

### C. Data collection and processing

Surface EMG of 12 muscles was recorded with bipolar electrodes (Delsys Bagnoli, Delsys Incorporated, Boston, MA) at 2048 Hz. This included six ventral muscles: *Rectus Abdominis* (RA) at the umbilicus level, *External Obliques* (EO), and *Internal Obliques* (IO) and six dorsal muscles: *Longissimus Thoracis pars Thoracis* (LTpT), *Longissimus Thoracis pars Lumborum* (LTpL), and *Iliocostalis* (IL). A reference to their exact placement can be found here [10].

MATLAB (2022a, The Mathworks, Natick, MA) was used for data processing and further analysis. For data processing, we used the same pipeline as Moya-Esteban et al. [10]. All signals were filtered using a second-order zero-lag Butterworth filter with different cut-off frequencies (*f*_*c*_). Linear EMG envelopes were obtained by consecutively bandpass filtering (*f*_*c*_ : 30 − 300 Hz), full-wave rectifying, and low-pass filtering (*f*_*c*_ : 6 Hz) of the recorded EMG signals. EMG linear envelopes were normalized using data from maximum voluntary contraction (MVC) recordings^4^. After filtering, all recorded signals were resampled at 50 Hz to manage the data size. After pre-processing, data from each lifting cycle were resampled to 1001 time points, representing the entire lifting phase from start (0%, just before the participant began reaching for the box, standing straight) to end (100%, just after the participant had returned to the initial standing position, standing straight).

### D. EMG reconstruction from a reduced sensor set

In this section, we explain our approach to reconstruct EMG channels from a limited number of sensors. We begin by discussing data splitting to obtain both a training and a test set. Subsequently, we describe how we extracted the synergy vectors from the calibration set, followed by the selection of optimal sensors. Finally, we demonstrate how we reconstructed the EMG for unmeasured channels.

Apriori, we evaluated the data from Moya-Esteban et al. and found that the ventral muscles (RA, EO, and IO) had less influence during lifting [10]. Thus, we perform the synergy analysis using only the six dorsal muscles.

Fig. 2 shows a flowchart of the sensor reduction method, where *m* − *k* muscles are reconstructed from *k* sensors for *n* samples. Here, *m* indicates the total number of included dorsal muscles, and *k* equals the number of synergies.

**Fig. 2.**
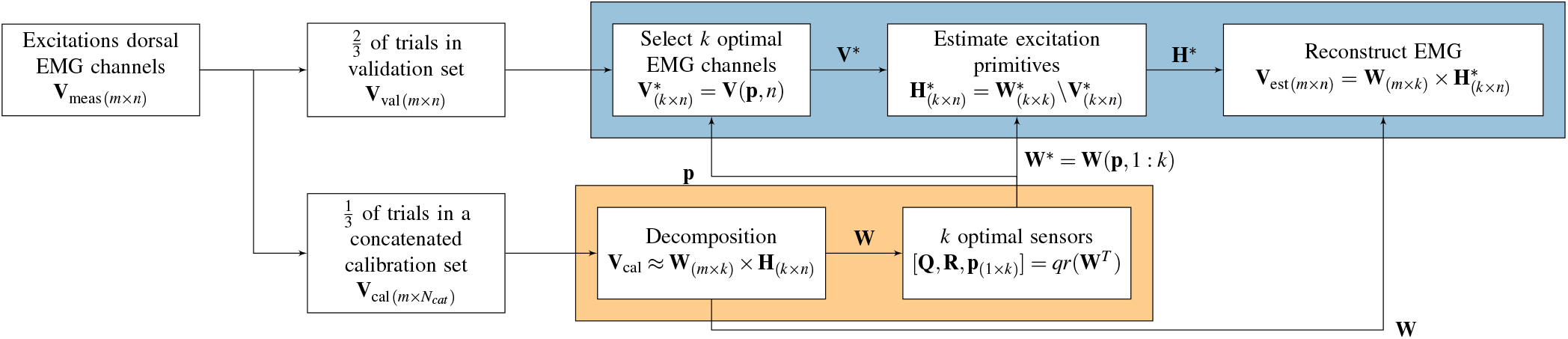
This flowchart outlines the inverse synergy-based workflow. The orange box shows the calibration steps, while the blue box indicates the reconstruction steps. We reconstructed *m* − *k* unmeasured EMG channels in **V**_est_ for *n* samples and *k* sensor locations using this approach for each participant. The first step involves dividing the EMG linear envelopes of *m* muscles (**V**_meas_) into a validation set (**V**_val_) and a calibration set (**V**_cal_). For each participant, **V**_cal_ consists of EMG signals of all *m* muscles, and selected calibration trials (with concatenated length *N*_*cat*_). The **V**_cal_ is the input of the NMF algorithm to obtain a time-invariant synergy vector matrix **W**_(*m*×*k*)_. From **W**_(*m*×*k*)_, *k* optimal sensor locations (indicated by **p**_(1×*k*)_) are chosen using a QR factorization algorithm, and a reduced synergy vector matrix 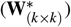 is created from their corresponding entries in **W**_(*m*×*k*)_. Only these *k* sensors from **V**_val_ are treated as measured (indicated by 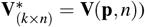). The time-variant excitation matrix 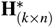 is estimated using a least-squares approximation (eq. 2). Finally, **V**_est_ is reconstructed from 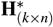 and the calibrated **W**_(*m*×*k*)_ (eq. 3).

#### 1) Data Splitting

Initially, trials were categorized randomly into a calibration dataset **V**_cal_ (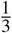 of the data) to obtain the synergy vectors and select optimal sensor locations, and a validation dataset **V**_val_ (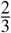 of the data) to evaluate the EMG reconstruction accuracy. So for each movement-weight combination, two out of six trials were used in the calibration dataset for ST and SQ, while one out of three trials was used for UT and BT. This ratio was chosen to ensure that a sufficient number of trials were included in the validation dataset to evaluate the reconstruction accuracy. All trials selected for the calibration dataset were concatenated (which results in length *N*_*cat*_ in 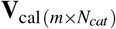.

#### 2) Synergy extraction from calibration data

In the first calibration step, we determined the time-invariant synergy vector matrix **W**_(*m*×*k*)_. To achieve this, we applied NMF to **V**_cal_ with *k* ranging from one to six, using MATLAB’s NNMF function:

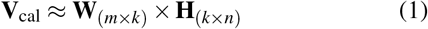

resulting in the synergy vector matrix **W**_(*m*×*k*)_, and the synergy excitation matrix **H**_(*k*×*n*)_. The identified **W**_(*m*×*k*)_ was then saved and later used for obtaining **V**_est_ across all combinations of movement and weight.

#### 3) Optimal sensors from calibration data

The second calibration step involved selecting the optimal locations for *k* sensors. The minimum number of sensors was determined by identifying the minimum number of synergies needed to explain VAF ≥ 0.90, as often used in literature [18], [20]–[22], [24], [25], also known as the threshold method [18]. Variability calculation is further explained in (II-E). Once the minimum number of sensors was established, the goal was to choose the most effective sensor locations from among the dorsal sensors. This process focused on creating a submatrix 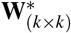 from the matrix **W**_(*m*×*k*)_. Here, 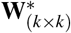 was used to estimate movement-dependent synergy excitations, 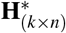. This estimation was based on the measured EMG signal of the most effective sensor locations 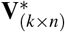 and involved solving a linear least squares problem given by:

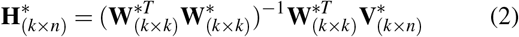

To improve reconstruction accuracy, it is important to maximize the absolute value of the determinant of the submatrix 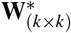, which represents its volume. The optimal sensor locations were determined by the rows of matrix **W**_(*m*×*k*)_ that contribute to the largest submatrix volume in 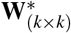. To find these rows, QR factorization with column pivoting of (**W**_(*m*×*k*)_)^*T*^ was used [26]. The optimal *k* sensor locations were found by the first *k* pivot locations in the pivot vector, i.e. **p**_(1:*k*)_. These pivots best sampled the submatrix 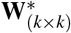.

#### 4) EMG reconstruction of the validation data

After obtaining the submatrix 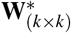, we could estimate the movement-dependent synergy excitation matrix 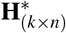 for each timestep, as illustrated in equation (2). The final step involved EMG reconstruction, expressed by:

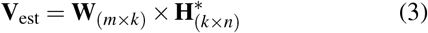

To assess the accuracy of **V**_est_ using the reduced sensor set, we applied this method to reconstruct all trials within the validation set **V**_val_.

### E. Data analysis

To find the minimum number of synergies *k* from the calibration set (**V**_cal_) that corresponds to the number of columns of matrix **W**_(*m*×*k*)_ from NMF, we used the VAF, defined as follows:

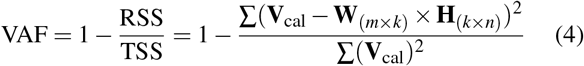

Here RSS and TSS are the residual and total sum of squares, respectively. The optimal number of decomposition columns in **W**_(*m*×*k*)_, which corresponds to the optimal number of sensors, was found when the median VAF exceeds 0.90 for a specific *k*. VAF values range from 0 (no explained variability) to 1 (perfect fit).

For model evaluation, we compared only reconstructed channels in **V**_est_ to their measured values in **V**_val_ (indicated as reference values). We used root mean squared errors (RMSE) to evaluate the average magnitude of prediction errors for *n* samples:

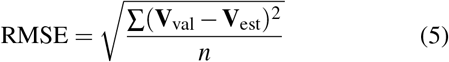

A lower value of RMSE indicates better model performance, with 0 indicating a perfect fit.

We used the coefficient of determination (*R*^2^) to evaluate the extent to which the variance of the model (**V**_est_) explains the variance of the reference (**V**_val_):

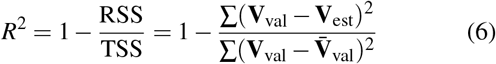

Note that while the definitions of VAF and *R*^2^ are similar, the TSS for VAF is concerning zero, whereas for *R*^2^, it is related to the mean 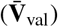 [18]. Unlike VAF, *R*^2^ can be negative, indicating poorer performance than a constant predictor using the mean (where, *R*^2^ = 0). The maximum value for *R*^2^ is 1.

## III. RESULTS

Table I shows that when the number of synergies increases from one to six, the median VAF of the reconstructed calibration data also increases, while the interquartile range (IQR) decreases. For *k* ≥ 2, we see a median VAF *>* 0.90. This indicates that subject-specific synergy matrices for *k* = 2 (i.e. **W**_(6×2)_ and 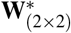) can be applied for EMG estimation.

**TABLE I.**
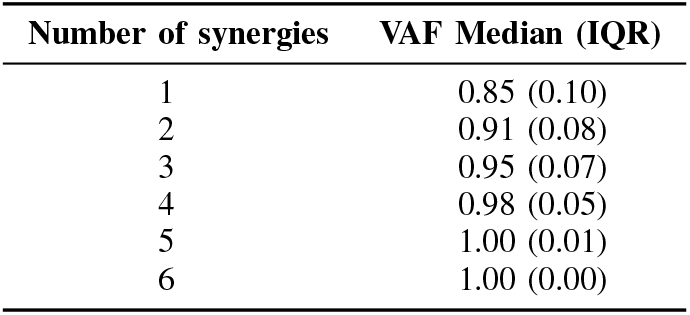
Reconstruction VAF of the NMF decomposed calibration data of all subjects. For ≥ 2 synergies, the median VAF *>* 0.90.

Table II shows where sensors were placed on all participants using the QR factorization algorithm. The LTpL muscles were chosen most frequently (16 times). Upon examining sides, the left side was favored more than the right. Further investigation revealed that, out of the two sensor locations, all participants except one had a bilateral sensor placement. This single exception had two left sensors, which accounts for the difference between the totals for the left and right sides.

**TABLE II.** The optimal sensor locations for all participants, per muscle (columns) and side (rows). Totals are provided in italics.

| Sensor location | LTpT | LTpL | IL | $\Sigma_{\text{Side}}$ |
| --- | --- | --- | --- | --- |
| Left | 3 | 9 | 1 | *13* |
| Right | 2 | 7 | 2 | *11* |
| $\Sigma_{\text{muscle}}$ | *5* | *16* | *3* | *24* |

Fig. 3 illustrates a representative participant. Reconstruction using the inverse synergy-based approach is demonstrated across several weights, including 0 kg (a), 7.5 kg (b), and 15 kg (c) for an SQ, and several lifting techniques, including an ST (d), UT (e), and BT (f) with 15 kg weight. The chosen sensor locations for this participant were LTpT left and IL right. The model scales EMG based on weight, and the temporal alignment of peaks in the reconstructed signal corresponds to those in the reference signal. However, differences exist between measured and reconstructed EMG; for instance, in (f), the model predicts a peak around 85% of the lifting phase, which was not measured.

**Fig. 3.**
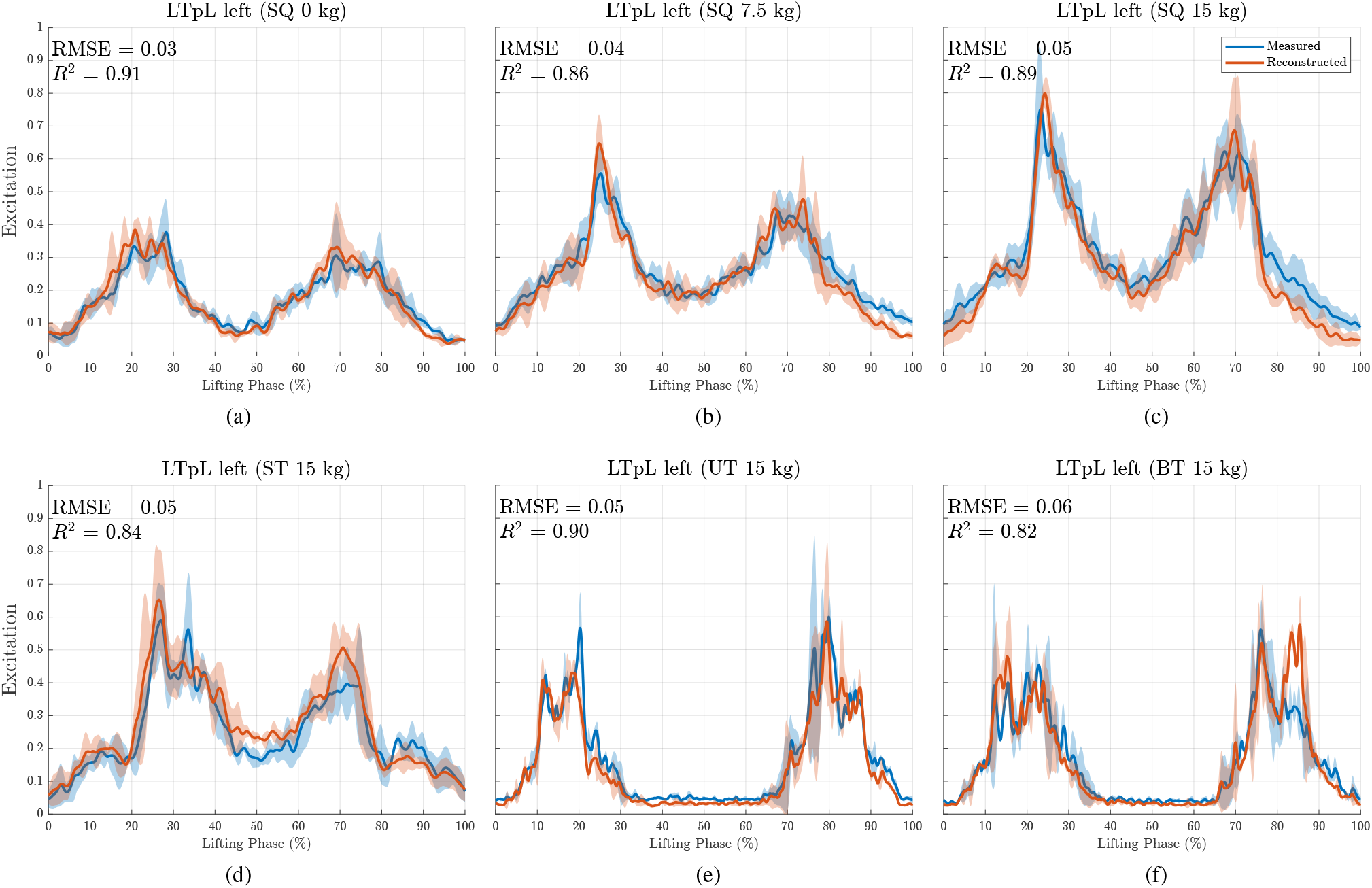
Measured and reconstructed mean EMG (solid line) and standard deviation (shaded region) of one participant (subject 6) for the left LTpL muscle (sensor locations are LTpT left and IL right). Plots a-c show the EMG during SQ liftings with increasing weight, respectively 0 kg (a), 7.5 kg (b), and 15 kg (c). Plots c-f show the EMG for lifting 15 kg using different lifting techniques, respectively SQ (c), ST (d), UT (e), and BT (f).

Table III shows median RMSE and *R*^2^ for reconstructed EMG for all movement and weight conditions. We observe median RMSE values ranging between 0.02 (ST 0 kg and UT 0 kg) and 0.04 (BT 15 kg, SQ 15 kg, ST 7.5 kg and ST 15 kg). Generally, the RMSE increases with higher weights. The median *R*^2^ values range between 0.70 (UT 0 kg) and 0.86 (SQ 15 kg). Similarly, we observe an increasing *R*^2^ with higher weights.

**TABLE III.** RMSE and *R*^2^ between **V***val* and **V***est* for reconstructed EMG. Median values (IQR) across all participants are shown.

|  | EMG<br>(reconstructed sensors only) |  |
| --- | --- | --- |
| | RMSE [-] | $R^2$ [-] |
| BT 0 kg | 0.03 (0.02) | 0.71 (0.41) |
| BT 7.5 kg | 0.03 (0.02) | 0.78 (0.31) |
| BT 15 kg | 0.04 (0.02) | 0.81 (0.20) |
| SQ 0 kg | 0.03 (0.02) | 0.78 (0.36) |
| SQ 7.5 kg | 0.03 (0.02) | 0.81 (0.16) |
| SQ 15 kg | 0.04 (0.02) | 0.86 (0.11) |
| ST 0 kg | 0.02 (0.02) | 0.76 (0.22) |
| ST 7.5 kg | 0.04 (0.02) | 0.74 (0.14) |
| ST 15 kg | 0.04 (0.02) | 0.83 (0.14) |
| UT 0 kg | 0.02 (0.01) | 0.70 (0.39) |
| UT 7.5 kg | 0.03 (0.01) | 0.81 (0.31) |
| UT 15 kg | 0.03 (0.02) | 0.83 (0.21) |

Fig. 4 shows the joint distribution of RMSE and *R*^2^ values. Some outliers have *R*^2^ *<* 0, especially for 0 kg and 7.5 kg. Additionally, we see more instances of 0 ≤ *R*^2^ ≤ 0.5 for 0 kg compared to 7.5 kg and 15 kg. In contrast, RMSE outliers are related to higher weights. To determine the presence of peaks in the measured EMG signal, we looked if the EMG values exceeded a threshold of 5% (of the maximum excitation) above the mean EMG signal value for a minimum of 10% of the lifting phase. Trials failing to surpass this threshold are indicated with a cross. We see that many outliers for *R*^2^ *<* 0 do not contain EMG peaks above this threshold.

**Fig. 4.**
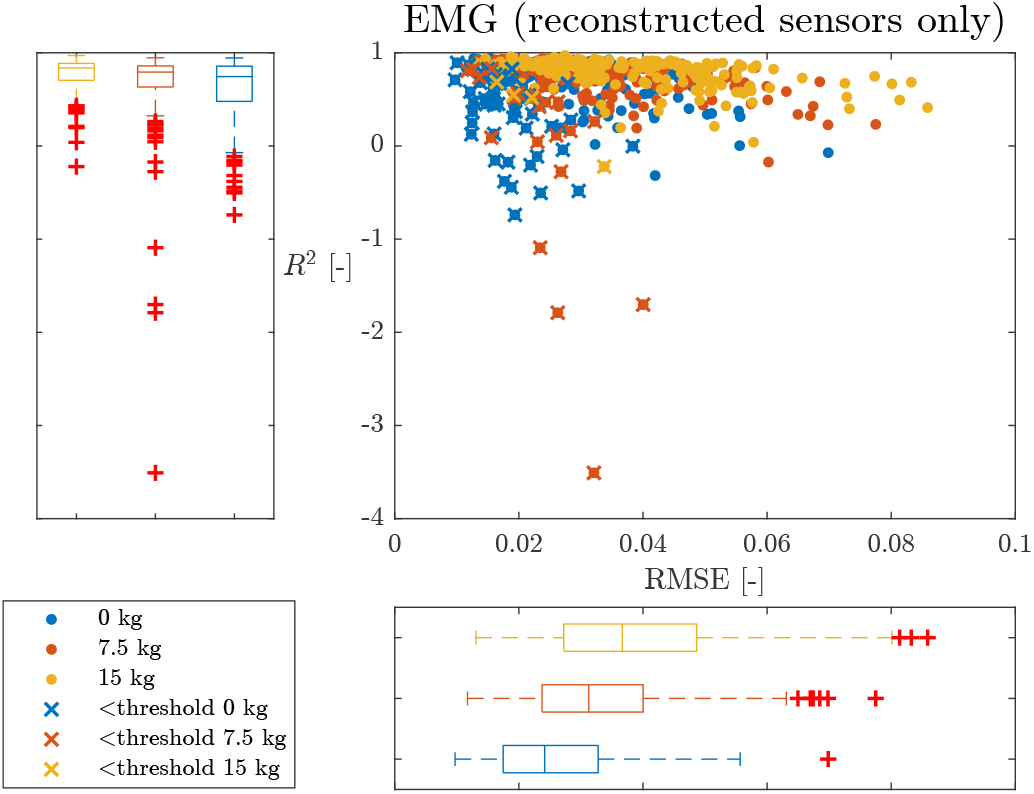
Boxplots and scatter plot showing the joint distribution of RMSE and *R*^2^ values for reconstructed muscles only (4 muscles×12 participants×12 conditions). Crosses indicate data points for trials where EMG excitation values do not exceed the threshold of 5% (of the maximum excitation) above the mean signal value for at least 10% of the lifting phase.

## IV. DISCUSSION

Our main aim was to improve the wearability of our trunk EMS model using an inverse synergy-based approach. This was assessed by testing the feasibility of reconstructing EMG signals of unmeasured muscles using a reduced sensor input. We evaluated our approach for occupational lifting tasks, including symmetric (SQ, ST) and asymmetric (UT, BT) techniques, and various weight conditions (0, 7.5, 15 kg).

The threshold method resulted in two optimal synergies that had a median VAF *>* 0.90. Obtaining the optimal number of synergies avoids too many synergies, which results in an overdetermined system and is undesirable for a minimal approach. Conversely, having too few synergies could result in loss of useful information. Our findings are consistent with Wang et al., [21] who also identified that two muscle synergy groups explain VAF *>* 0.90. Their study encompassed various movements, including flexion/extension, lateral bending, axial rotation, sitting, and standing. Notably, these movements, except sitting, are relevant to symmetric and asymmetric box lifting. Sedaghat-Nejad et al. [20] identified a set of four synergies. Differences between their findings and ours could arise from including ventral muscles in their synergy analysis. This may necessitate a greater number of synergies to explain synergy excitation variances. Similar reasons could explain the differences with Tan et al. [19], who identified three synergies but included the biceps brachii and gluteus maximus muscles in their analysis.

We observed that also using one synergy achieves a reasonably good median VAF (0.85). This is due to the similarity in EMG excitation patterns for left- and right-sided dorsal muscles during symmetric lifting movements. However, during asymmetric lifting, EMG peaks may be missed due to expected peaks at different instances of the lifting phase for left and right-sided muscles. The identified muscle synergies align with the functional anatomy, because the back behaves as one unit during symmetric lifting tasks, and two units during asymmetric tasks.

To examine sensor locations, we used QR pivots to maximize the absolute value of the determinant of the reduced synergy vector matrix, **W**^*^. This method aims to increment the volume of **W**^*^ constructed from the pivoted columns of **W**, preventing an ill-conditioned system. When using two sensors, selecting sensors corresponding to the highest column values in the first and second columns of **W** yields the same outcome as employing QR pivots. As the number of sensors increases, choosing the most optimal ones becomes less evident [25]. In these situations, QR pivots could provide a simple and effective strategy [26].

QR factorization identified optimal sensor locations on either side of the back. LTpL was commonly chosen as the optimal sensor location, except for one participant, who had two optimal sensors identified on the left side. The reason for this difference must be explored further.

The results indicate that using two sensors allows for the reconstruction of the EMG of four unmeasured muscles, with RMSE values ranging from 0.02 for ST 0 kg and UT 0 kg to 0.04 for BT 15 kg, SQ 15 kg, ST 7.5 kg and ST 15 kg. Overall, we observed low RMSE for all lifting techniques and box weights. Median RMSE values increased for increased box weights. This can be explained because lifting heavier weights is characterized by peaks in the EMG excitation signal. Time delays between the reconstructed and reference peaks may cause an increased RMSE and reconstruction errors in peak height.

Median *R*^2^ values range from 0.70 (UT 0 kg) to 0.86 (SQ 15 kg). However, we observed significant outliers that occasionally fall below zero (fig. 4). These outliers were predominantly associated with median (7.5 kg) and low (0 kg) box weights. Notably, low *R*^2^ values did not exhibit a direct correlation with high RMSE. We hypothesize that baseline differences caused by noise or constant tonic muscle activity may account for these variations, as mentioned by [27]. This could have a large influence on the *R*^2^, since it is sensitive to these errors, as they significantly impact the differences in the mean of the reconstructed and reference signals. Variations in the signal-to-noise ratio (SNR) between reconstructed and measured channels could contribute. Specifically, if a reconstructed signal from a measured EMG exhibits a high SNR in comparison to a reference signal with a low SNR, it may lead to a low *R*^2^.

The results of this study indicate that two dorsal sensors can accurately estimate the EMG of four unmeasured dorsal sensors with low RMSE. This suggests that we can derive joint moments and compression forces through an EMS using only two sensors as input, instead of requiring to map all muscle activity of the back [10]. Placing EMG sensors in the correct anatomical locations on the skin is a tedious process. Therefore, sensor reduction is essential to improve the wearability and applicability of an EMS in occupational settings. This facilitates ambulatory biomechanical risk assessment in occupational settings by monitoring compression forces to evaluate the effect of ergonomic interventions [14], [28]. Sensor reduction also improves the usability of an EMS as a human-machine interface (HMI) to control an active back-support exoskeleton [10], [13].

The study has a few limitations. In order to identify subject-specific synergies, we had to measure all muscles during calibration, and assumed that the identified synergy vectors did not change for the subsequent sessions. Future research should focus on identifying population-wide synergies preferably from diverse groups [29]. This could standardize the calibration procedure using population-based optimal sensor locations, eliminating the need to measure all muscles during calibration for each participant.

Furthermore, we did not include the influence of muscle fatigue in our analysis. Fatigue could induce motor unit (MU) rotation (alternating activity between MUs) in the trunk extensor muscles [30], [31]. Therefore, synergy vectors could alter during lifting tasks when fatigue induces MU rotation, influencing the reconstruction accuracy of estimated EMG channels. Online techniques could constantly update these vectors reflecting changes in the muscle activity. Finally, as a next step, we need to validate if our reduced approach can inform a trunk EMS to provide reliable estimates of lumbosacral joint moments and compression forces.

## Footnotes

3 From hereon, these conditions are used to indicate the weight conditions.

4 From hereon, EMG refers to these normalized EMG linear envelopes, unless stated otherwise.

## Notes

* This work was supported by the European Union’s Horizon 2020 RIA (Grant No. 871237; SOPHIA), and Horizon 2024 RIA (Grant No. 101120408; SWAG).

### Competing Interest Statement

The authors have declared no competing interest.

